# Reconstructing physiological oxygen gradients reveals the role of hypoxia in colon epithelial organization

**DOI:** 10.64898/2025.12.16.694730

**Authors:** Jiaquan Yu, Chuyi Chen, Felicia H. Rodriguez, Gianfranco L. Yee, Isabella Y. Zeng, Kevin Yang, Elizabeth K. Benitez, Karuna Ganesh, Scott R. Manalis

## Abstract

Oxygen gradients organize tissue architecture and metabolism^1,2^, yet their precise spatial profiles and mechanistic roles remain poorly understood because both *in vivo* measurement and *in vitro* control are technically challenging^3,4^. Here, we quantify the oxygen landscape of the mammalian intestine using microscale sensors, revealing a steep luminal–basal gradient of approximately 10-60 µM mm^−1^ that collapses under antibiotic perturbation. We then recreate this physiological range *ex vivo* with a submerged chemostat microfluidic platform that fixes the oxygen boundary condition by coupling an oxygen-permeable PDMS chip to an external scavenger reservoir and integrating embedded optical sensors for real-time readout. This architecture suppresses ambient oxygen ingress and sustains programmable gradients of 10-20 µM mm^−1^ across three-dimensional colorectal cancer organoid cultures while remaining compatible with live imaging and endpoint retrieval. The platform bridges quantitative *in vivo* oxygen mapping with controlled *ex vivo* modeling, establishing a generalizable approach to interrogate how spatial oxygen dynamics govern epithelial organization and disease progression.

## Introduction

Oxygen gradients are fundamental to tissue physiology, guiding development, metabolism, and adaptation5. In the intestine, oxygen declines sharply from the arterioles in the submucosa to the mucosal surface, shaping epithelial polarity and microbial ecology^6,7^. Although absolute oxygen levels have been measured in several organs^8^, the spatial derivatives that define gradient steepness and continuity remain poorly characterized2. This lack of quantitative gradient information obscures whether disruptions in oxygen distribution directly modulate intestinal physiology and cancer progression.

Three-dimensional human organoids derived from healthy donors or patient tumors provide powerful models for epithelial biology and disease. Yet these cultures typically experience uniform oxygenation far exceeding physiological levels, because ambient oxygen readily diffuses through polymers and media^9–11^. Efforts to impose hypoxia, using materials with low oxygen diffusivity and gas-controlled incubators, have achieved partial success but are limited by instability, lack of quantitative control, and incompatibility with live imaging^12,13^. A platform that can stably reproduce *in vivo* gradients while enabling direct measurement and visualization would close this critical gap.

Here, we introduce a scavenger-submerged microfluidic chemostat that overcomes these limitations. By immersing a PDMS chip in a heated oxygen-scavenger bath and a confined microfluidic oxygen supply, the device maintains a precise oxygen boundary at the chip level. This setup eliminates ambient leakage, permits placement of embedded optical oxygen sensors within 200 µm of the culture, enables real-time oxygen readout, and supports multi-day 3D organoid culture under physiologically relevant gradients, without specialized incubators. We then show that this system reconstructs intestinal oxygen landscapes across defined microfluidic geometries, linking *in vivo* measurements to tunable *ex vivo* environments.

## Results

To define the native oxygen landscape of the intestine, we performed microsensor-based measurements in the large intestine of euthanized C57BL/6 mice under normoxic conditions. After dissection, a needle-encapsulated trace-range electrode was advanced along the colonic lumen. In wildtype mice, luminal oxygen remained <2 µM distal to the cecum and throughout the colon, whereas antibiotic-treated mice, characterized by epithelial inflammation and depletion of commensal microbiota, exhibited significantly elevated luminal oxygen with a mean of 40 µM (**Fig. 1a**). Together with published values (**Extended Data Table 1**), these data indicate that a substantial luminal-to-basal oxygen difference is a hallmark of colon physiology and collapses when the homeostasis is disrupted.

**Fig. 1.**
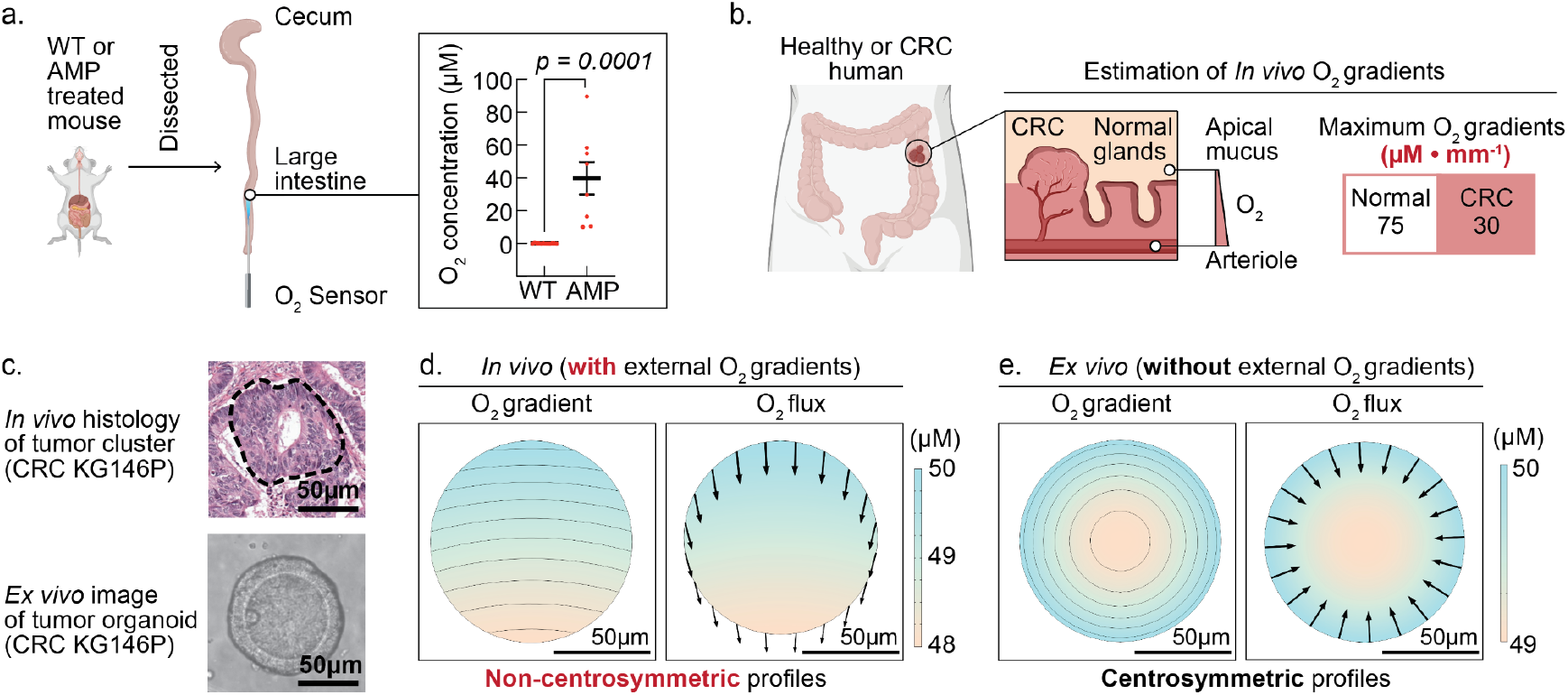
*ex vivo* organoid culture exhibits a different oxygen profile than *in vivo* tissue. **a**. *in vivo* measurement of oxygen levels in mouse intestine; **b**. Estimated maximum oxygen gradient ranges for healthy and CRC humans; **c**. Histology of CRC patient KG146P showing ductal architecture cancer *in vivo*, shown in black circles. 3D grown *ex vivo* organoids from KG146P recapitulates the *in vivo*-like cystic architectures; **d**. Simulated *in vivo* oxygen concentration profile showing non-centrosymmetric oxygen profiles; **e**. Simulated oxygen concentration profile of *ex vivo* grown KG146P organoids showing that oxygen profiles differ significantly from *in vivo* to *ex vivo* conditions, even when maintained at stable ambient hypoxia of 50 µM.

To determine the actual oxygen gradient and enable human–mouse comparison, we analyzed histological sections from a CRC patient (KG146P) and previous reports (**Extended Data Table 1**). By measuring the distance from the mucous surface to adjacent vasculature and integrating published intratumoral oxygen measurements, we estimated that maximum *in vivo* human CRC cancer gradient is 30 µM mm^−1^ (**Fig. 1b**). Although *ex vivo* organoids recapitulate key aspects of tumor architecture (**Fig. 1c**), their oxygen profiles diverge markedly. Specifically, finite-element simulations (COMSOL) indicate that external gradients above 10 µM mm^−1^, when imposed across epithelial structures, dominate intrinsic oxygen consumption, driving directional oxygen fluxes that are non-centrosymmetric (**Fig. 1d,e**). Together, these results underscore that the key feature of 30 µM mm^−1^ oxygen gradient *in vivo* has yet to be reconstructed and tested *ex vivo*.

Building on the physiological oxygen gradients quantified *in vivo*, we next sought to recreate oxygen profiles *ex vivo* using a controllable microfluidic system. To build such a chemostat culture platform, we leveraged the oxygen permeability of polydimethylsiloxane (PDMS) to develop a submerged microfluidic culture chip capable of generating physiologically relevant oxygen gradients (**Fig. 2a**). This approach directly addresses the limitations of previous methods and provides two distinct advantages.

**Fig. 2.**
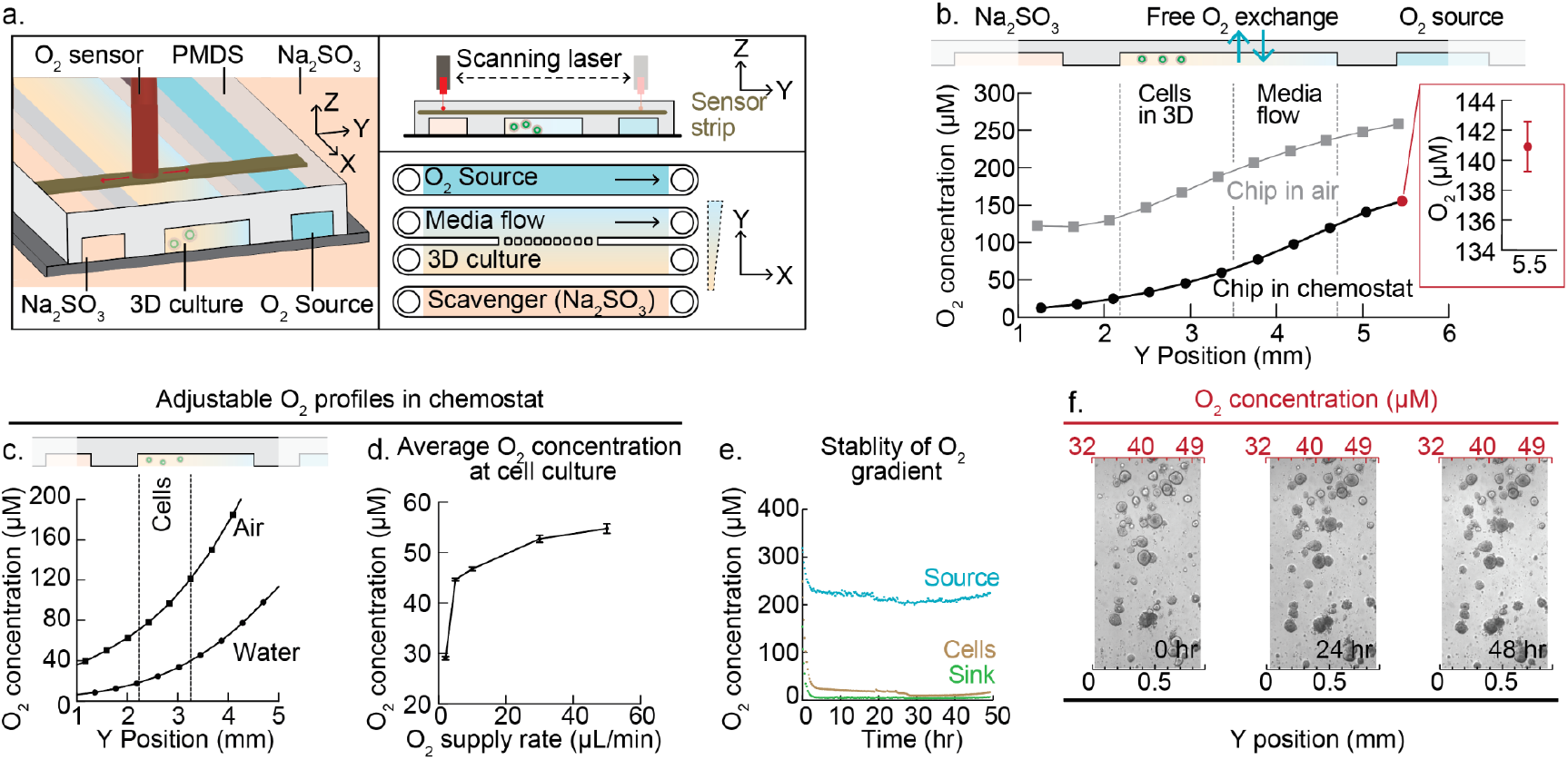
Chemostat system to maintain colorectal cancer organoids with stable trans-organoid oxygen gradients. **a**. Schematic of the chemostat comprising a four-channel PDMS chip submerged under sodium sulfite oxygen scavenger, integrated with an oxygen-scanning sensor and time-lapse microscopy to profile stable oxygen gradients across 3D cultures; **b**. Measured oxygen profiles demonstrating that scavenger submergence is required to achieve physiologically relevant intestinal hypoxia; **c and d**. Tunable oxygen gradients (20–120 µM) across the cell culture region achieved by varying (**c**) oxygen source and (**d**) flow rate of oxygen source (air saturated water); **e**. Continuous real-time tracking of oxygen concentration over 50 hours showing gradient stability using air-saturated water as oxygen source at a flow rate of 2 µl/min.; **f**. Combined oxygen scanning and time-lapse microscopic imaging, enabling quantitative mapping of oxygen levels and gradients across individual organoids from patient KG146P.

First, submerging the chip in a heated, oxygen-scavenger–filled tank prevents ambient oxygen exchange (**Extended Data Note 1**). This configuration eliminates the need for hypoxia incubators^14^ while allowing an oxygen sensor system and bright-field imaging to be fully integrated on-chip (**Extended Data Fig. 1a**). Taking advantage of the accessibility of the submerged design, we embedded a customized, high-sensitivity oxygen sensor strip within the PDMS channel and positioned less than 200 µm away from the ECM and organoids (**Extended Data Fig. 1b**). A fiber-optic probe continuously scans the sensor strip to read localized oxygen concentration in real time (**Fig. 2a Extended Data Fig. 2a**), generating complete oxygen profiles every seven minutes (**Extended Data Fig. 2b**).

Second, by placing the scavenger outside the PDMS chip and introducing on-chip oxygen source and scavenger sink channels, we achieved stable, physiologically relevant oxygen levels and gradients (**Fig. 2b**–e). Notably, both empirical measurements (**Fig. 2b**) and COMSOL simulations (**Extended Data Note 2**) revealed that the scavenger-submerged configuration is essential to achieve physiological hypoxia. Without the submerged scavenger, oxygen concentrations exceed 100 µM within 500 µm of the on-chip sink (**Extended Data Note 2**), a limitation previously reported for devices lacking ambient isolation^13^.

Confining the oxygen source within the chip while surrounding it with a liquid scavenger as an external sink enables precise control over oxygen gradients through three key design features. First, the simplicity of the design allows oxygen gradients to be generated in a wide variety of microfluidic configurations, provided that an oxygen source channel is incorporated^15,16^. For example, we fabricated four- and five-channel chips with varying culture-channel dimensions and source-to-culture distances, each producing distinct oxygen gradient profiles (**Extended Data Fig. 3b,c**). Second, separating the oxygen source channel from the cell-culture channels with a PDMS barrier enables gas exchange without direct liquid contact^17^. This configuration allows the use of either mixed gas or air-saturated water as the oxygen source (**Fig. 2c**) and permits independent flow-rate control in the source channel without imposing shear stress on the cultured cells (**Fig. 2d Extended Data Fig. 3a**). Finally, water flow rates can be controlled with much greater precision than airflow, and the slower diffusion rate of oxygen in water allows finer control over hypoxic conditions^18^. Hypoxia establishes in the tank within one minute (**Extended Data Note 1**), and adjustable oxygen gradients in the range of 30 µM to 120 µM form across the culture region within one hour (**Fig. 2b,c**; **Extended Data Note 2**).

**Fig. 3.**
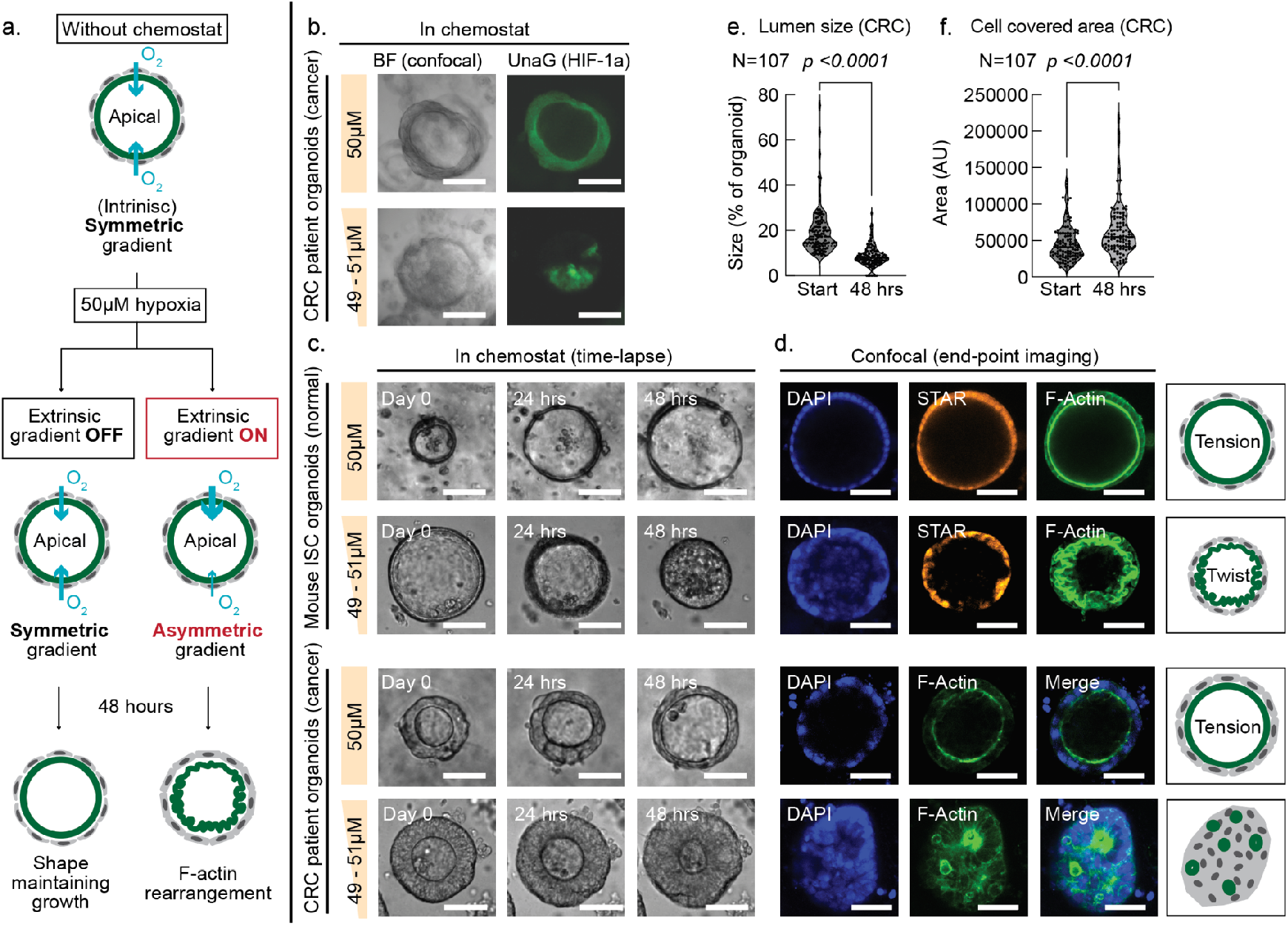
Oxygen gradients govern colorectal organoid architecture. **a**. Schematic illustrating how an imposed oxygen gradient induces polarity reversal and F-actin rearrangement in patient-derived CRC organoids; **b**. Fluorescence images of patient-derived CRC organoids expressing the UnaG hypoxia reporter under uniform hypoxia or an imposed oxygen gradient, showing even UnaG signal distribution under uniform conditions and its concentration on the hypoxic side under a gradient; **c**. Bright-field images from the chemostat chip showing outward (cystic) growth under uniform hypoxia versus inward (compact) growth under an oxygen gradient; **d**. Confocal micrographs depicting organoid morphology under uniform hypoxia compared to hypoxic-gradient culture; **e**. Quantification of inward growth under an oxygen gradient as a reduction in lumen size over 48 h; **f**. Quantification of cell-covered area under an oxygen gradient over 48 h; **e,f**. P values from paired two-tailed t-tests with Welch’s correction; **b.c.d**. Scale bars are 100µm.

Because the sodium sulfite–based scavenger reacts with oxygen faster than oxygen can diffuse into the liquid, it effectively prevents oxygen from reaching the culture region (**Extended Data Note 1**). Stability was maintained until the scavenger was fully neutralized, a process mitigated by replacing the scavenger every two days (**Fig. 2e Extended Data Note 1**). This approach not only produces robust and reproducible oxygen gradients but also supports long-term studies of oxygen-dependent biological processes in 3D organoid cultures.

The ability to precisely map oxygen gradients across a 3D organoid culture in the chemostat enables direct correlation between oxygen levels and organoid behavior (**Extended Data Fig. 4a,b**). Using bright-field imaging, we spatially annotated individual organoids according to their position within the gradient (**Fig. 2f**), quantifying both absolute oxygen levels (in µM) and gradient steepness (in µM mm^−1^). The system ensures stable oxygen conditions with less than 5 % variation (**Extended Data Fig. 4c**) across the 10 mm culture region (X-axis) and yields more than ten organoids per oxygen level per chip. Together, these features provide a practical and generalizable platform for reconstructing physiological oxygen gradients *ex vivo*, linking quantitative *in vivo* measurements to tunable, stable microenvironments for mechanistic studies of 3D tissue organization.

**Fig. 4.**
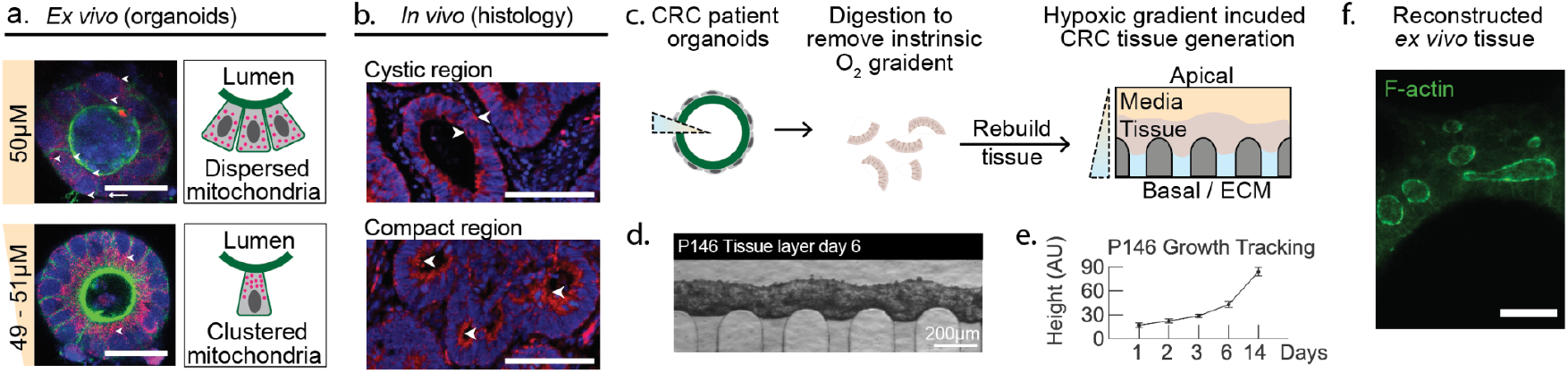
Hypoxia-induced tissue compaction enables reconstruction of *ex vivo* CRC tissue with histological heterogeneity. **a**. Patient-derived CRC organoids cultured under chemostat-generated oxygen gradients show mitochondrial clustering (red) on the apical side of nuclei (blue). **b**. Immunofluorescence of patient tumor KG146P tissue sections stained for TOMM20 (red) and DAPI (blue) reveals regions with dispersed mitochondria (top) and regions with apically polarized mitochondria (bottom), echoing the *ex vivo* phenotype. **c**. Schematic illustrating reconstruction of CRC tissue under imposed oxygen gradients, following digestion of organoids to remove intrinsic oxygen profiles. **d**. Bright-field image showing continuous epithelial tissue formed after 6 days of chemostat culture. **e**. Real-time imaging enables longitudinal tracking of tissue growth in the chemostat. **f**. F-actin staining highlights heterogeneous lumen sizes within reconstructed tissue, resembling patient histology. **a,b,f**. Scale bars, 100 µm. d, scale bar, 200 µm.

In colorectal cancer (CRC), tumor progression is closely aligned to the oxygen gradient along the luminal-basal axis of the intestinal epithelium^19^. CRC typically originates from intestinal stem cells at the base of colonic crypts, initially forming polyps that protrude into the hypoxic intestinal lumen^20,21^. Such hypoxia-associated polyps can cause bleeding and obstruction, but are benign and can be endoscopically resected, preventing progression to cancer^22^. As polyps progress to cancer, they invade basally, accessing oxygen-carrying blood vessels, and from there, disseminate to distant sites^23^. Migration along this apical-basal oxygen gradient is therefore a key prognostic and predictive feature of CRC, yet the specific role of the oxygen gradient in driving polarity reversal in CRC cells has not been previously studied due to the lack of physiological *ex vivo* culture systems^24^. Like the normal colon, CRC and patient-derived CRC organoids self-organize into polarized structures with F-actin-bounded apical lumen and an ECM-facing basal layer^25^. A primary tumor consists of multiple such glandular epithelial “organoid”-like structures encapsulated in ECM and stroma, and organized along the apical-basal oxygen gradient of the intestine (**Extended Data Fig. 5**). Mechanical disruption, such as physical dissection or transitioning from ECM to suspension culture, can induce reorganization, altering the apical-basal orientation^25^. This polarity coexists with an intrinsic oxygen gradient: higher oxygen levels on the basal (ECM and capillary facing) side and hypoxic conditions on the apical (luminal) side^26^.

Leveraging our chemostat platform’s ability to generate stable and precisely tunable oxygen gradients, we investigated a key question: how does growth in an oxygen gradient, in comparison with hypoxia per se, influence CRC polarity and morphology? Studying this distinction is challenging because cellular oxygen consumption naturally generates oxygen gradients (luminal hypoxia/basal high oxygen from media) as organoids grow, complicating efforts to isolate and analyze these effects. Using our platform, we established a controlled environment to systematically test how hypoxia versus oxygen gradients influence CRC architecture (**Extended Data Fig. 6a,b**).

Since there are no reported measurements of oxygen gradients in 3D *ex vivo* organoid cultures, we first sought to estimate how external oxygen gradients influence the intrinsic luminal/basal gradients established by cellular oxygen consumption. Using COMSOL simulations, we modeled an external, *in vivo*-like oxygen gradient that disrupted the centrosymmetric oxygen profile, creating a non-centrosymmetric trans-organoid gradient (**Fig. 3a Extended Data Fig. 6a**). Initially, we simulated oxygen gradients generated solely by intrinsic cellular consumption. Due to the absence of reported oxygen consumption values for organoids under hypoxic conditions, we relied on literature-derived estimates to model a range of possible oxygen gradients. In all scenarios, intrinsic oxygen profiles remained centrosymmetric, with a hypoxic core exceeding 48 µM. Next, we incorporated an external oxygen gradient resembling physiological *in vivo* conditions, such as those achievable using the chemostat system (**Extended Data Fig. 6b**). Simulations of a 200 µm diameter organoid with a 50 µm shell thickness, corresponding approximately to a single-cell layer, revealed that external gradients on the order of 10-20 µM mm^−1^ significantly disrupted the intrinsic oxygen consumption pattern. Notably, the external gradient reversed the direction of oxygen flux at the hypoxic side, driving oxygen from the organoid core toward the luminal surface. These findings highlight the critical role of externally applied gradients in overriding the centrosymmetric gradient driven by cellular consumption and reshaping oxygen profiles within tissues.

Numerous studies have reported how hypoxia influences the polarization and differentiation of CRC, particularly through pathways regulated by HIF-1α^27^. We thus hypothesized that the intrinsic centrosymmetric oxygen profile plays a pivotal role in governing organoid growth and shape through classical hypoxia-driven mechanisms involving HIF-1^28^. To examine the influence of hypoxia, we transduced patient-derived CRC organoid KG146P^29^ with a UnaG reporter which encodes 5 copies of the hypoxia response element directing the expression of a destabilized oxygen-independent fluorescent protein (dUnaG) ^30,31^. UnaG transfected organoids displayed negligible autofluorescence under ambient oxygen conditions but reliably exhibit green fluorescence in hypoxic culture. When cultured under uniform hypoxic conditions at 50µM oxygen, the KG146P organoids uniformly expressed UnaG, representing global HIF-1 activation (**Fig 3b**). However, when exposed to a mild oxygen gradient averaging 50 µM, even with a minimal oxygen difference (<2 µM) across the organoid, we observed polarized UnaG expression, with higher signal intensity localized to the hypoxic side.

Interestingly, this disruption of the centrosymmetric oxygen profile was associated with inward growth of the organoids, with gradual filling of the organoid lumens (**Fig 3c**). This inward growth process commenced after ∼12 hours of oxygen gradient culture and progressed until the lumen was no longer visible. Both mouse intestinal stem cell-derived and primary patient-derived CRC organoids displayed intraluminal growth (**Fig. 3c**). In patient-derived KG146P CRC organoids exposed to oxygen levels below 50 µM, 91.6% of individual organoids showed a significant reduction in lumen size over 48 hours (**Fig. 3e**) accompanied by an increase in cell-covered area (**Fig. 3f**), yielding a net expansion in overall organoid volume that mirrors the luminal filling, proliferation, and biomass accumulation characteristic of tumors *in vivo*^32^. Organoids can be maintained in chemostat for over a week with continued growth, allowing assessment of reversibility of the luminal filling phenotype. Remarkably, reverting oxygen gradient back to normoxia fully restored the original cystic morphology of KG146P CRC organoids within 48 hours (**Extended Data Fig. 7**). These findings demonstrate that the external oxygen gradient can act to dynamically and reversibly sculpt epithelial architecture, inducing luminal filling and epithelial overgrowth that are characteristic hallmarks of tumorigenesis.

We next asked whether oxygen gradient-induced morphological changes are associated with cytoskeletal rearrangements and alterations in epithelial cell polarity and stemness. The chemostat chips, with the PDMS layer mounted on a glass coverslip, are compatible with conventional confocal microscopy (**Extended Data Fig. 1b**). Following two days of culture under a 30 µM mm^−1^ oxygen gradient, organoids were fixed in situ to prevent readaptation, and their apical-basal organization was analyzed using confocal microscopy. We first asked whether induction of an oxygen gradient plays a role in organizing the stem-differentiation axis of the intestinal epithelium by transducing mouse intestinal organoids with the Stem Cell ASCL2 Reporter (STAR)^33^. In mouse intestinal stem cell organoids expressing the STAR reporter, oxygen gradient exposure led to a spatial location-specific loss of stem cell gene expression in the most hypoxic part of the gradient where the oxygen flux was reversed by the external gradient (**Fig. 3d**). This observation suggests that the oxygen gradient likely plays a role in inducing the differentiation gradient observed in the normal intestine, where stem cells normally reside at the (most oxygenated) base of intestinal crypts. Phalloidin staining of F-Actin confirmed that both mouse normal and human cancer organoids grown under uniform 50 µM hypoxia exhibited typical apical-basal polarization, characterized by a dense, smooth apical (luminal) F-actin layer. However, when exposed to an external oxygen gradient, this actin layer appeared twisted, disorganized, and redistributed across both the apical and basal sides of the organoids (**Fig. 3d**).

In contrast, CRC primary tumor organoids, which are epigenetically restricted to an intestinal stem cell state^34^, retained STAR expression regardless of the oxygen gradient. However, approximately 20% of the CRC organoids developed a compact, no-lumen morphology during prolonged culture, in which the apical F-actin lining was disrupted and redistributed internally (**Fig. 3d**). Since active migration is associated with mitochondrial localization between the cell nucleus and F-actin in the direction of migration^35^, we stained CRC patient organoids with Mitotracker FM Deep Red, DAPI, and F-Actin to visualize relative location of mitochondria, nucleus and polarity, respectively. Staining of CRC primary organoids fixed ^36^ hours after exposure to the oxygen gradient revealed mitochondria concentrated around the actin structures, in between the nucleus and apical lining actin, which is consistent with the directionality of active migration toward the actin-aligned filling lumen (**Fig. 4a**). Importantly, the compact morphology and apically concentrated mitochondria induced by chemostat-generated oxygen gradients closely mirror the histology of the cognate tumor tissue from patient KG146P. Within the same tumor, glandular structures containing regions of evenly distributed mitochondria and regions of apically polarized mitochondria towards the lumen are detectable (**Fig. 4b**). These disease-relevant morphological features, which are absent in organoids grown under standard normoxia but can be induced by culture in oxygen gradients in the chemostat, implicate the physiological oxygen gradient of the intestine as an important driver of the luminal filling, migration and dense growth that are characteristic of cancer progression in the clinic. To further extend the chemostat platform from single-organoid phenotypes to tissue-level reconstruction, we dissociated CRC organoids and seeded them against the ECM interface to generate continuous epithelial layers under controlled oxygen gradients (**Fig. 4c**). Within six days, these fragments compacted into a cohesive tissue that could be tracked in real time by bright-field imaging, enabling longitudinal analysis of growth dynamics in the chemostat (**Fig. 4d,e**). Fixed tissues stained for F-actin revealed heterogeneous lumen sizes within the continuous layer (**Fig. 4f**), closely resembling the duct-like variability observed in patient tumor histology (**Extended Data Fig. 5**).

## Discussion

Contemporary microfluidic hypoxia platforms are often custom engineered, requiring bespoke integration of oxygen sensors, fluidics and imaging components^36–38^. To address these limitations, we established a set of design principles, developed a modular integration framework, and assembled a catalog of commercially available components that enable the generation of precise oxygen gradients in PDMS-based microfluidic systems. This approach provides a practical and reproducible pathway for enhancing chip-based platforms with tunable oxygen control.

Our modular system, combining an external scavenger reservoir with embedded PDMS-integrated sensors, can be readily adapted for applications ranging from organotypic culture and cell sorting to chemical synthesis and multi-omic assays. From a biological perspective, we show that spatial oxygen gradients, rather than uniform hypoxia, are key regulators of colorectal cancer organoid morphology and polarity associated with invasive tumor behavior. These findings highlight the importance of re-evaluating the role of hypoxia in cancer biology, developmental processes, and tissue physiology, specifically distinguishing the effects of oxygen gradient dynamics from those of absolute oxygen concentration.

## Supporting information

All extended data

## Acknowledgements

We thank Dr. Arnold Levine, Dr. Dan Littman, and Dr. Matthew Vander Heiden for helpful discussions and Annie Chen for facilitating device fabrication. This work was supported by Stand Up To Cancer (SU2C) Convergence Program 3.1416, D. K. Ludwig Fund for Cancer Research, and the MIT Center for Precision Cancer Medicine. This work was also supported in part by the Koch Institute Support (core) Grant P30-CA014051 from the National Cancer Institute.

