## Supplementary material for "Reconstructing physiological oxygen gradients reveals the role of hypoxia in colon epithelial organization": All extended data

Affiliations
[1]Koch Institute for Integrative Cancer Research, Massachusetts Institute of Technology, 500 Main St building 76, Cambridge, MA 02139, USA
[2]Department of Biological Engineering, Massachusetts Institute of Technology, 21 Ames St #56-651, Cambridge, MA 02139, USA
[3]Molecular Pharmacology Program, Sloan Kettering Institute, Memorial Sloan Kettering Cancer Center, New York, NY 10065, USA
[4]Weill Cornell/Rockefeller/Sloan Kettering Tri-Institutional MD-PhD Program, New York, NY 10065, USA
5Department of Medicine, Memorial Sloan Kettering Cancer Center, New York, New York 10065, USA
[6]Broad Institute of Harvard and MIT, 415 Main St, Cambridge, MA 02142, USA
[7]Department of Mechanical Engineering, Massachusetts Institute of Technology, 33 Massachusetts Ave, Cambridge, MA 02139, USA

* Authors contributed equally

### Extended Data Table 1

#### 1.1 Overview: Using Oxygen Scavenger to Enable Hypoxia Culture

| Condition | Length scale (mm) | Reference | Location | $O_2$ Concentration Range (µM) | Reference | Maximum $O_2$ Gradient (µM $mm^{-1}$) |
|---|---|---|---|---|---|---|
| **Human (Healthy)** | 0.7 to 1.4 | 1 - 3 | Arteriole | 5 - 60 | 4, 5 | 75 |
| | | | Lumen | 6 - 14 | 5, 6 | |
| **Human (CRC)** | 1.4 to 3.9 (Measured) | KG146P | Arteriole | 14 - 42 | 6, 7 | 30 |
| | | | Lumen | 4 - 15 | 8 | |
| **Mouse (Healthy)** | 0.2 to 0.5 (Measured) | 9 | Arteriole | 56 - 60 | 4, 6 | 250 |
| | | | Lumen | 10 - 22 | 4 | |
| **Mouse (Disease)** | 0.2 to 0.3 (Measured) | 10 | Arteriole | 25 - 75 | 4 | 315 |
| | | | Lumen | 12 - 40 | 4 | |

*Maximum oxygen gradient was defined as the largest arteriole–lumen oxygen difference divided by the shorter length scale, with values rounded to the nearest 5-unit increment.

### Extended Data Note 1

#### 1.1 Overview: Using Oxygen Scavenger to Enable Hypoxia Culture

To establish and maintain a hypoxic environment for organoid culture, we designed a submerged tank system filled with a sodium sulfite ($Na_2SO_3$) solution, which functions as a chemical oxygen scavenger. The tank was constructed as a cuboid with internal dimensions of 8 × 11 × 8 $cm^3$, yielding a total volume of approximately 700 mL. The sodium sulfite actively reacts with dissolved molecular oxygen, limiting ambient oxygen diffusion into the surrounding medium and the embedded microfluidic chip. The relevant reaction proceeds via:

$$2\ Na_2SO_3 + O_2 \rightarrow 2\ Na_2SO_4.$$

We performed matched COMSOL simulations and experimental measurements to characterize two key phases of this oxygen-quenching process. In the initial phase, addition of sodium sulfite to oxygen-saturated water rapidly depletes dissolved oxygen, creating a uniformly hypoxic aqueous environment. Over subsequent days, oxygen levels remain near zero as the scavenger continues to neutralize oxygen diffusing in from the ambient air. Our simulation and empirical results consistently demonstrate that, as long as an adequate concentration of scavengers is maintained, the system sustains hypoxia throughout the culture period. Our findings are corroborated by numerical predictions, experimental data, and previously reported values from the literature, collectively validating the use of this submerged scavenger design for mutil-day hypoxic culture.

#### 1.2 Governing Equation and Parameters

Since there is no convective flow in the tank, the profile of oxygen concentration $C$ governed by the following diffusion-reaction equation:

$$\frac{\partial C}{\partial t} = D\nabla^2 C + R \qquad (1)$$

For the initial phase of oxygen quenching, the reaction term $R_o$ is pseudo first order [11] as:

$$R_o = -k_o C \qquad (2)$$

And for the subsequent scavenger quenching phase:

$$R_s = -k_s C_s^{0.65} \qquad (3)$$

$D$ is the oxygen diffusion coefficient in sodium sulfite solution, $k_o$ and $k_s$ are the reaction rate constants representing oxygen consumption by the sodium sulfite, $C_s$ is the sodium sulfite concentration, and $t$ is the time. The equation for R is determined by multiple studies demonstrating that the rate of the uncatalyzed oxidation of sodium sulfite in aqueous solution is independent of the dissolved oxygen concentration, indicating a reaction order of zero with respect to oxygen. The same studies report that the reaction rate depends on the sodium sulfite concentration, with the reaction experimentally determined to be approximately $0.65^{th}$ order [12,13]. The critical parameters used in the simulation are shown in **Table 1** below:

| Parameter | Value | Basis for Selection |
|---|---|---|
| Microfluidic chip position | The center of the microfluidic chip is located at point P with coordinates of (4 cm, 5.5 cm, 1 cm) within the tank | The position aligns with the information in **Extended Data Fig. 1a**. |

| | | |
|---|---|---|
| Oxygen diffusion coefficient in sodium sulfite | $\sim 2.75\times 10^{-9}\ m^{2}/s$ | Oxygen diffusion in sodium sulfite solution is approximated to that in water[14] |
| Oxygen solubility in water | $210\ \mu mol/L$ | Approximate saturation concentration in sodium sulfite solution at around physiological temperature [15] |
| Oxygen concentration in air | $20.95\%$<br>$(\sim 8.56\times 10^{-3}\ mol/L)$ | Oxygen concentration in air at room temperature calculated by the Ideal Gas Law; used as the ambient boundary condition |
| Initial sodium sulfite concentration | $4.26\ \ g/L$<br>$(\sim 33.8\times 10^{-3} mol/L)$ | Reflects the experimental setup. The sodium sulfite concentration was intentionally set at this high level to minimize the need for solution replacement and prevent disturbance to long-term organoid culture |
| Pseudo first order sodium sulfite–oxygen reaction rate constant, $k_o$ | $0.048\ \ s^{-1}$ ,$0.19\ s^{-1}$ | Various values considered based on literature [11] |
| Sodium sulfite–oxygen reaction rate constant, $k_s$ | $3\times 10^{-5} mM^{0.35}/s$,<br>$5.77\times 10^{-5} M^{0.35}/s$,<br>$1.23\times 10^{-5} M^{0.35}/s$ | Various values considered based on literature[12,13] |

**Table 1 | Constants used to simulate scavenger reaction.**

We assumed that at t = 0, the concentration of oxygen in the tank is $10^{-3}\ mol/L$ (saturated) everywhere, and at y = 0 cm and y = 8 cm, the concentration is always $2.1\times 10^{-4}\ mol/L$. Since the tank is air permeable, we assumed that oxygen diffuses into the tank freely.

### 1.3 Numerical Model Setup

#### 1.3.1 Hypoxia can be established in chemostat within a minute

We modeled the system in COMSOL and solved for the oxygen concentration at point P over time, and found that the oxygen level falls below detection (to ~ 0μM) after 39 seconds using the best-fit reaction rate constants, corresponding to the oxygen sensor data obtained from the Chemostat that indicates complete oxygen quenching within 35-40 seconds. (**Extended Data Note 1 Fig. 1**).

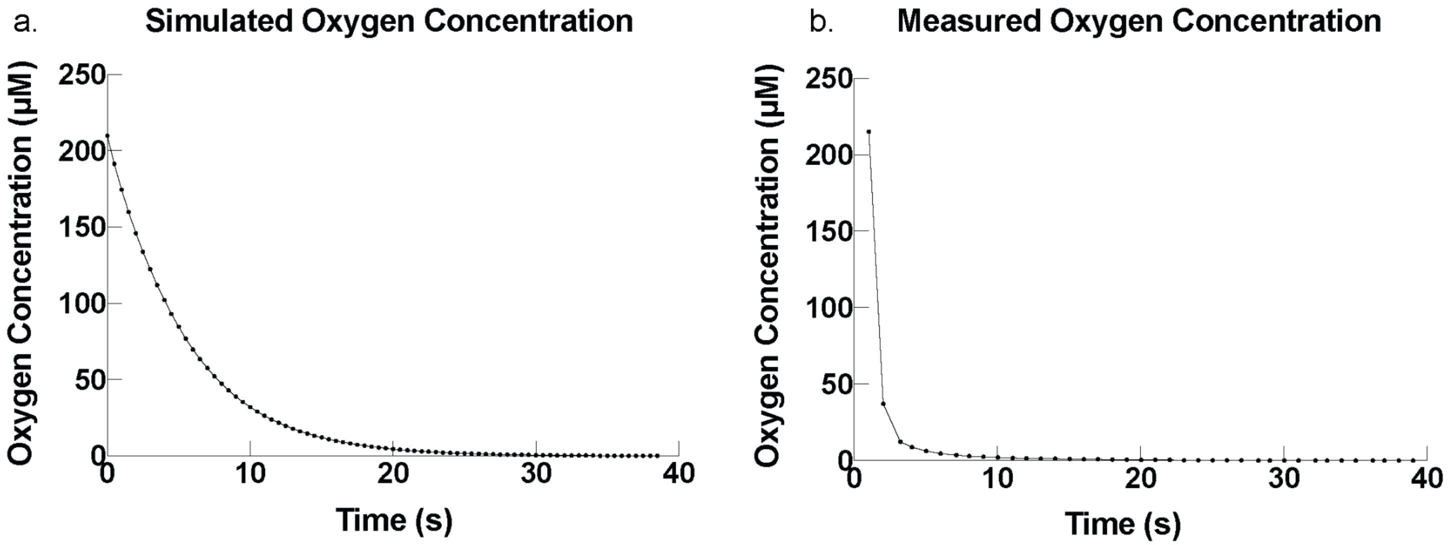


**Extended Data Note 1 Fig. 1 | Hypoxia in the chemostat can be established rapidly. a.** COMSOL simulation yields zero oxygen level at the center of the microfluidic chip (x=4 cm, y=5.5 cm, z=1 cm) within 39 seconds using the oxygen scavenger tank. **b.** The simulation is verified by the oxygen profile obtained using the oxygen sensor and the chemostat system, upper bounding the oxygen quenching time to 40 seconds. The first 20 seconds were used for adding the sodium sulfite into the solution, and temporarily stirring the solution to achieve a uniform scavenger concentration. The time needed in the empirical experiment for establishing a uniform concentration likely contributed to the delay of oxygen concentration drop and caused a sharper slope.

#### 1.3.2 Hypoxia can be maintained without addition of scavenger for >3 days

The corresponding COMSOL simulation was used to estimate the quenching time of the oxygen scavenger. All parameters and tank geometry were as previously described. Additional considerations include:

- A 1 cm-thick plastic tank wall was modeled in COMSOL. However, its impact on the quenching profiles is negligible due to the high oxygen diffusivity.
- At the surface of the tank, the oxygen concentration boundary condition was set to $210\ \mu M$, representing 20.95% of oxygen's solubility in water in the absence of reaction. This adjustment reflects the dominance of the rapid reaction over diffusion, which maintains the oxygen concentration at a significantly lower level than its equilibrium solubility.

The rate constant $k_s = 1.23\times10^{-5} M^{0.35}/s$ provided the best agreement with empirical data, yielding a scavenger quenching time of 100-110 hours. In the first 48 hours, the scavenger concentration remains above 6 μM. For approximately 4 days, the concentration of oxygen scavenger stays positive and the oxygen level remains at zero **(Extended Data Note 1 Fig. 2a)**, indicating that hypoxic conditions are stably sustained throughout this period. The simulation aligns with the oxygen concentration profile from the chemostat **(Extended Data Note 1 Fig. 2b)**.

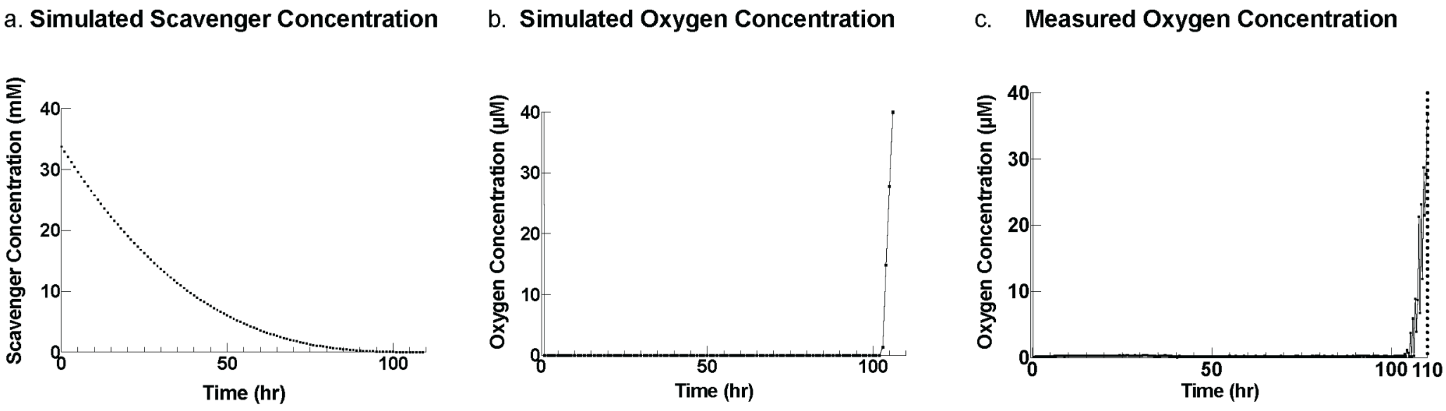


**Extended Data Note 1 Fig. 2 | Hypoxia in the chemostat can be established quickly and maintained over days. a&b.** The sodium sulfite concentration (averaged) in the tank shows that a positive amount of scavenger is consuming oxygen for more than 96 hours (4 days), resulting in complete oxygen depletion for 102 hours, and a subsequent oxygen level increase. **b.** Chemostat data agrees with the numerical analysis, showing a zero oxygen level within the first 100 hours, and a steep oxygen concentration rebound around the $100^{th}$-$110^{th}$ hours, aligning with the simulated time point for the scavenger quenching.

### Extended Data Note 2
### Modeling Oxygen Transport and Gradients in the Microfluidic Chip

To investigate oxygen concentration distribution within the microfluidic chip, a three-dimensional model was developed using finite element method (FEM)-based computational modeling software COMSOL Multiphysics (Burlington, MA, USA) [16]. The simulation domain consists of a PDMS block (15 x 25 x 4 mm³) housing the channels measuring 0.25 mm in height and 1.5 mm in width, bonded to a glass substrate at the bottom (**Extended Data Note 2 Fig. 1a**). The flow within channels was modeled using the "Laminar Flow" interface. The "inlet" conditions were applied to channel inlets with specified flow rates, and the "outlet" conditions were applied to the ends of the channels, with the pressure set to zero. The boundary conditions were set to "no slip" wall at the bottom, top, and sides of the channels. The flow within the channels was solved by a "stationary" solver. The "Transport of Diluted Species in Porous Media" interface was used to simulate oxygen diffusion through the PDMS, governed by Eqn. (1) (**Extended Data Note 1**). The oxygen concentrations in both the oxygen source and scavenger channels were assumed to be constant by applying a fixed value using a "Concentration" node. The oxygen concentration at the PDMS surface was assumed to be zero, as the chip was fully submerged in an excess of scavenger solution. The oxygen scavenging reaction in the scavenger channel was defined by specifying a reaction rate of 0.0577 mol/(m³·s) in the "Reaction" node. "Inflow" boundary conditions with specified flow rates were applied at the channel inlets, while "outflow" conditions were applied at the outlets. With these settings, oxygen transport within the PDMS and channels was simulated using a stationary solver.

A numerical model was employed to simulate oxygen profiles within the chemostat microfluidic chip under varying detector positions and oxygen supply conditions (**Extended Data Fig. 3b & c**), providing a computational reference for interpreting experimental measurements [17].

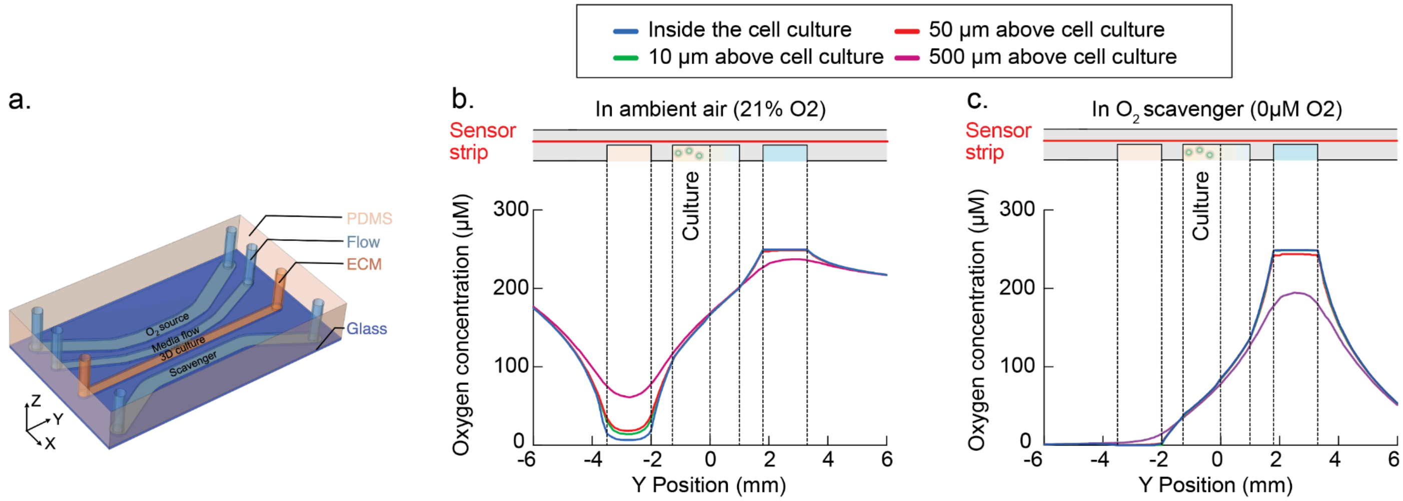


**Extended Data Note 2 Fig. 1| COMSOL simulation demonstrating that scavenger-submerged chips are essential to achieve physiologically relevant hypoxia. a.** Multi-channel microfluidic PDMS chip modeled in the COMSOL simulation. **b.** Simulated oxygen concentration across the entire PDMS chip exposed to ambient air. Oxygen levels within the cell culture region spanning ~100 µM to ~180 µM. **c.** Simulated oxygen concentration when the chip is submerged under sodium sulfite oxygen scavenger, where cell culture region spans ~30 µM to ~80 µM. **b&c.** Both simulations assume a 20 µL/min water flow as the oxygen source.

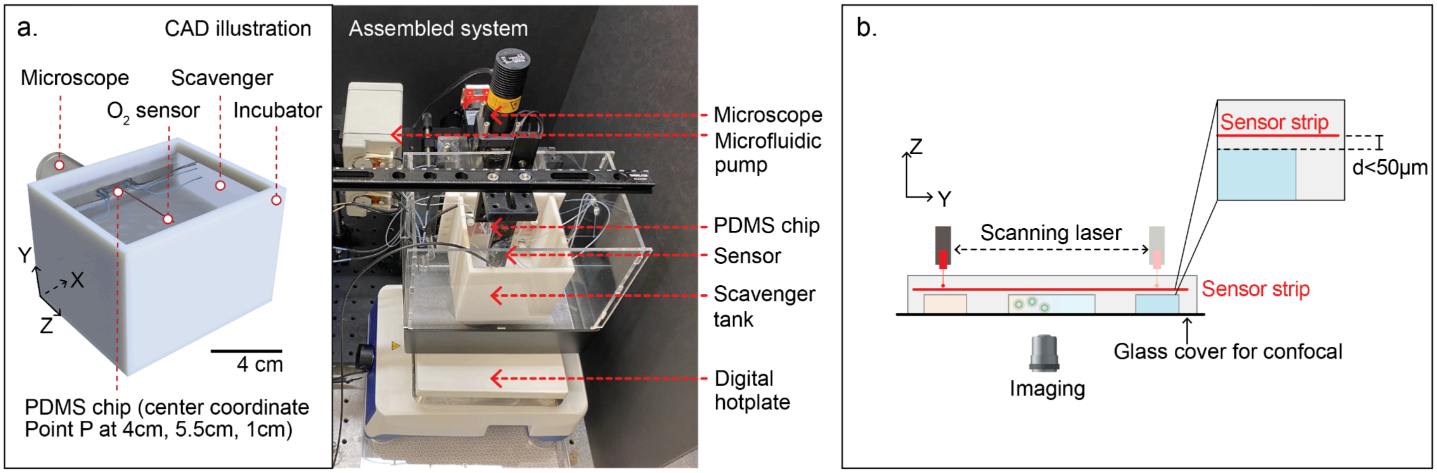


**Extended Data Fig. 1| Chemostat system enables coupled real-time live-cell imaging and adjustable hypoxia profiling. a.** Photograph of the experimental setup, showing the xyz-motorized stage, fiber-optic oxygen sensor, microfluidic pump, PDMS culture chip, imaging optics and temperature controller. **b.** Schematic of chemostat chip, illustrating back-side confocal access and PDMS-side oxygen scanning via a sensor strip embedded during PDMS casting less than 200μm from the cell culture for in situ oxygen measurement.

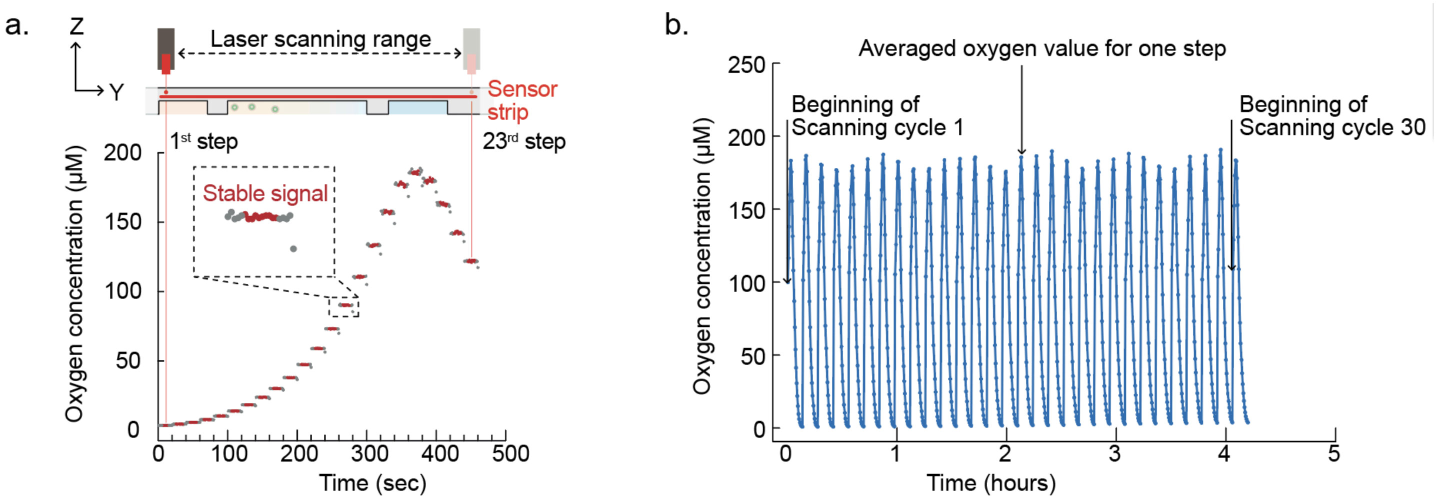


**Extended Data Fig. 2| Continuous *in situ* profiling of oxygen gradients. a.** Raw data from the fiber-optic sensor measured across 23 positions of the chemostat chip (step size 1 mm). At each position, the sensor dwells for 20 s, generating 20 measurements; the middle 10 stabilized readings are averaged to determine the local oxygen concentration. **b.** Sequential full-chip scans performed every 7 min produce cyclic oxygen profiles, demonstrating sustained gradient stability.

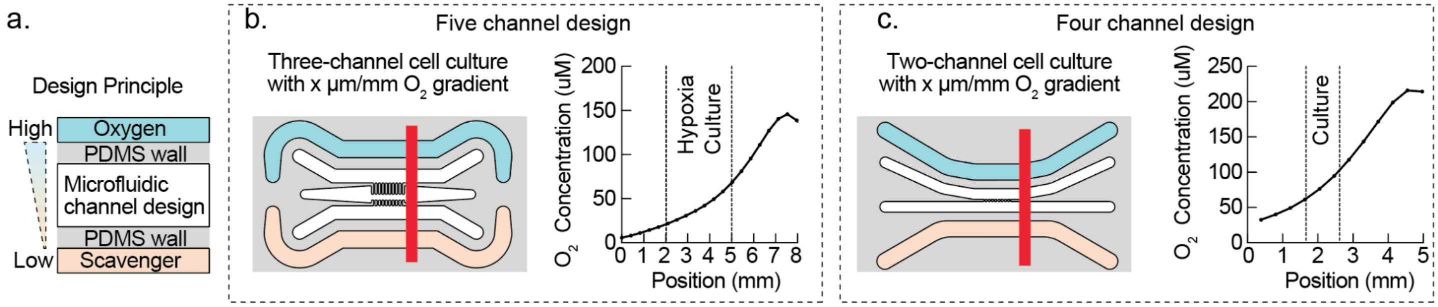


**Extended Data Fig. 3| Chemostat chip design principle enables generation of oxygen gradients in diverse microfluidic devices. a.** Schematic of the design principle: submerging the PDMS chip under sodium sulfite scavenger creates a hypoxic condition, requiring only a single oxygen-input channel and allowing straightforward integration with existing channel-based microfluidic systems. Oxygen and scavenger channels are physically separated from the cell culture channels with PDMS barrier walls to allow flow rate in the oxygen source channel to be independently controlled without inducing shear stress on the cultured cells; **b.** Five-channel configuration with three central culture channels, a common multiculture layout used in droplet generation, organ-on-chip and cell-assay applications, readily adapted to chemostat hypoxia; **c.** Four-channel design used in this study, wherein reduced spacing between the oxygen source and sink channels produces a steeper gradient compared to the five-channel layout.

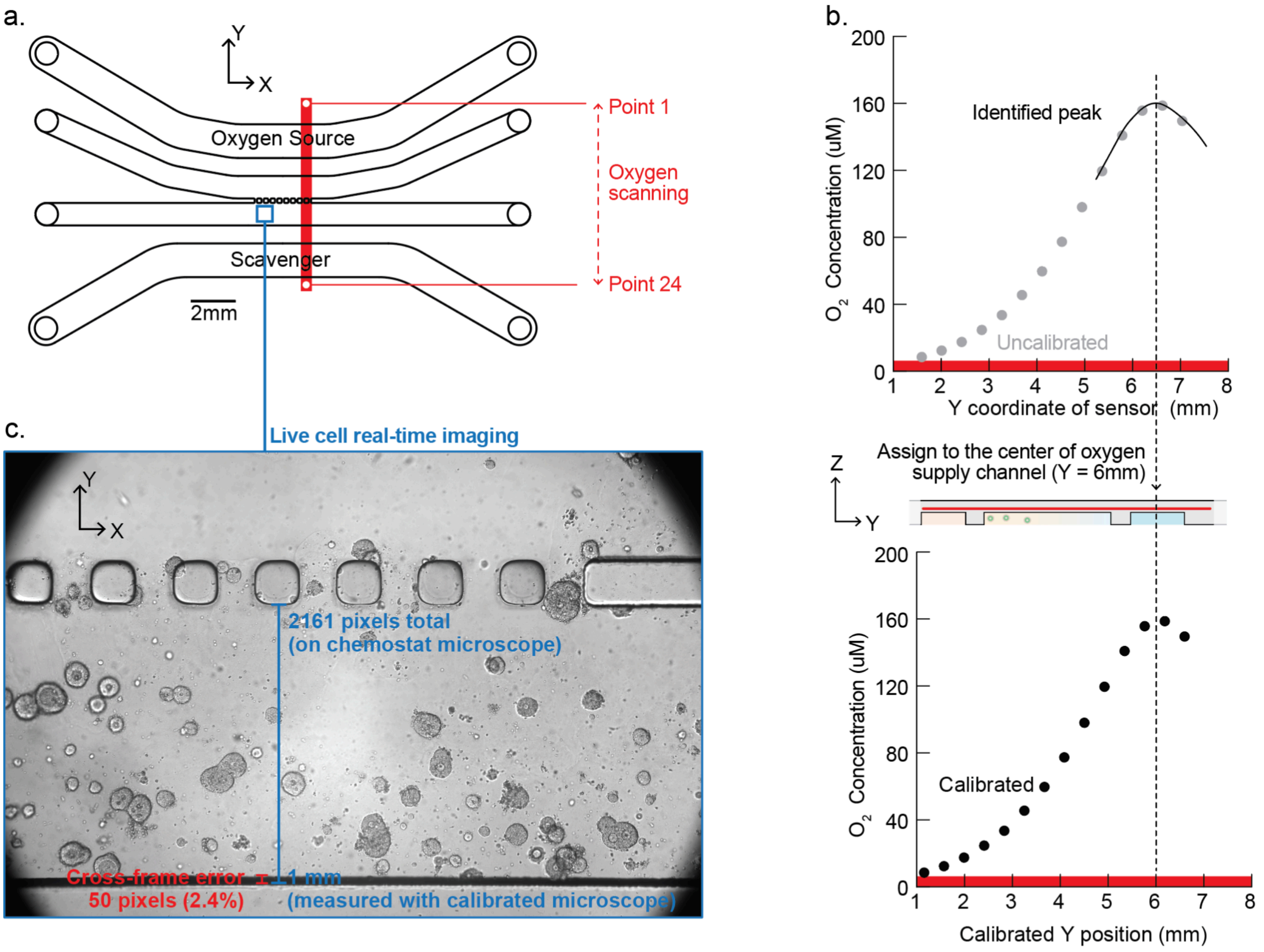


**Extended Data Fig. 4| Chemostat enables precise alignment of oxygen gradients with real-time live-cell microscopy. a.** Overlay of in situ oxygen measurements and live-cell imaging on the chemostat chip. **b.** MATLAB routines detect the peak oxygen level, corresponding to the center of the oxygen-supply channel, to map sensor coordinates (uncalibrated) onto chip coordinates (calibrated). **c.** Bright-field images provide intrinsic landmarks for direct registration to chip coordinates and correlation with the oxygen gradient; focus drift across frames introduces <5% alignment error (2.4% in this example).

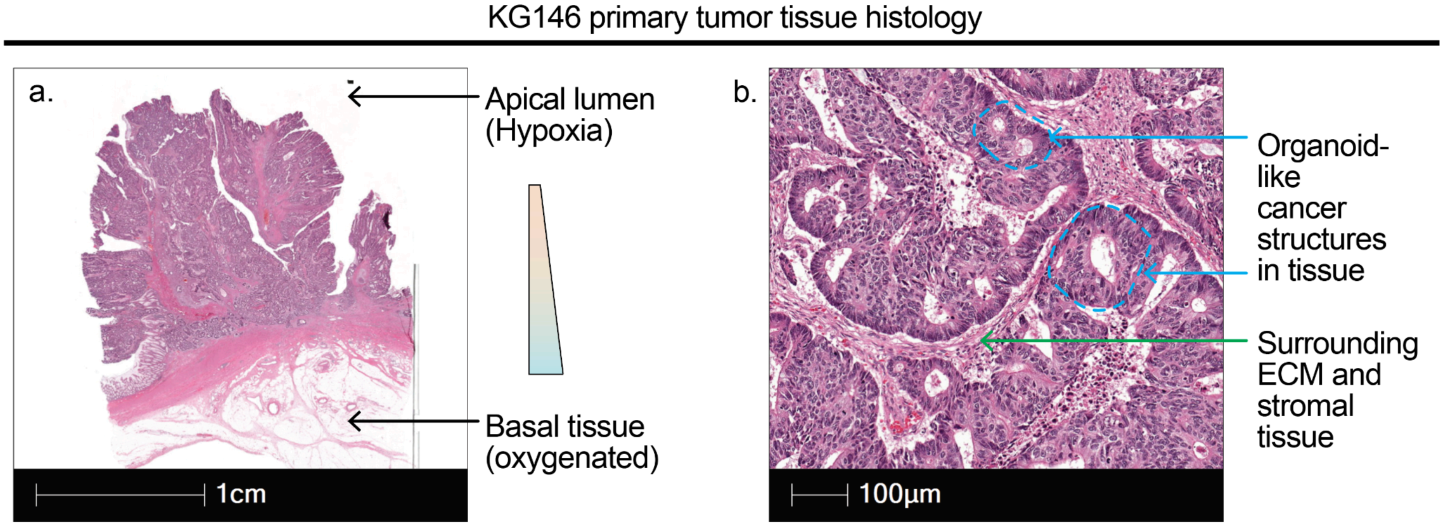


**Extended Data Fig. 5 | Histological organization of patient KG 146 primary colorectal tumor. a.** Low-magnification H&E-stained section illustrating the apical–basal orientation of the tumor *in situ*, aligned with the intestinal axis. **b.** High-magnification image revealing multiple glandular epithelial structures resembling organoids, each encapsulated within extracellular matrix and surrounded by stromal tissue.

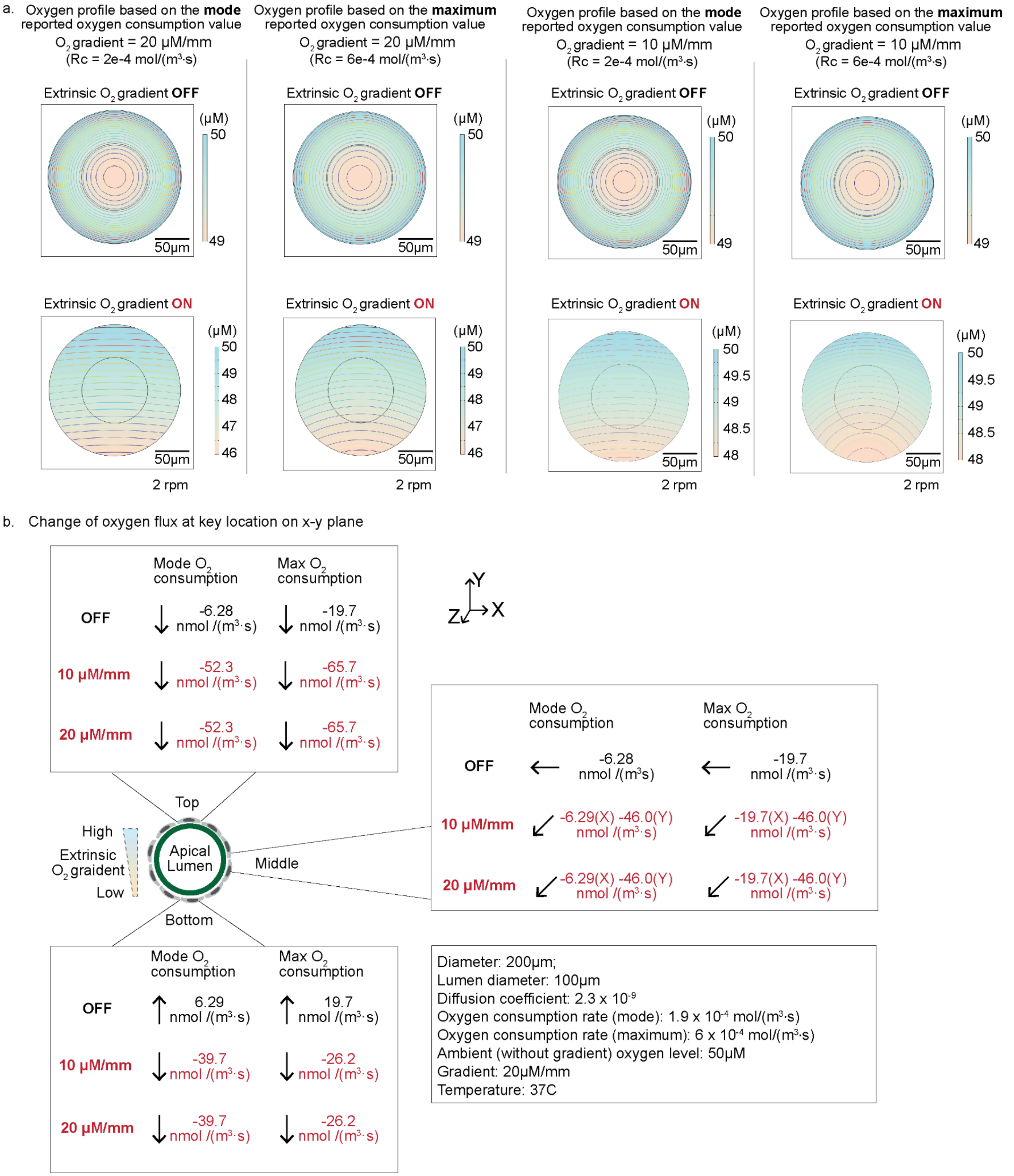


**Extended data Fig. 6| Oxygen profile of cultured organoid. a.** COMSOL simulation of oxygen gradient within organoids cultured with or without extrinsic oxygen gradient, using mode and the maximum of reported oxygen consumption of cells reported in literature. **b.** Calculated change of oxygen flux at key locations of each organoid, according to COMSOL simulation.

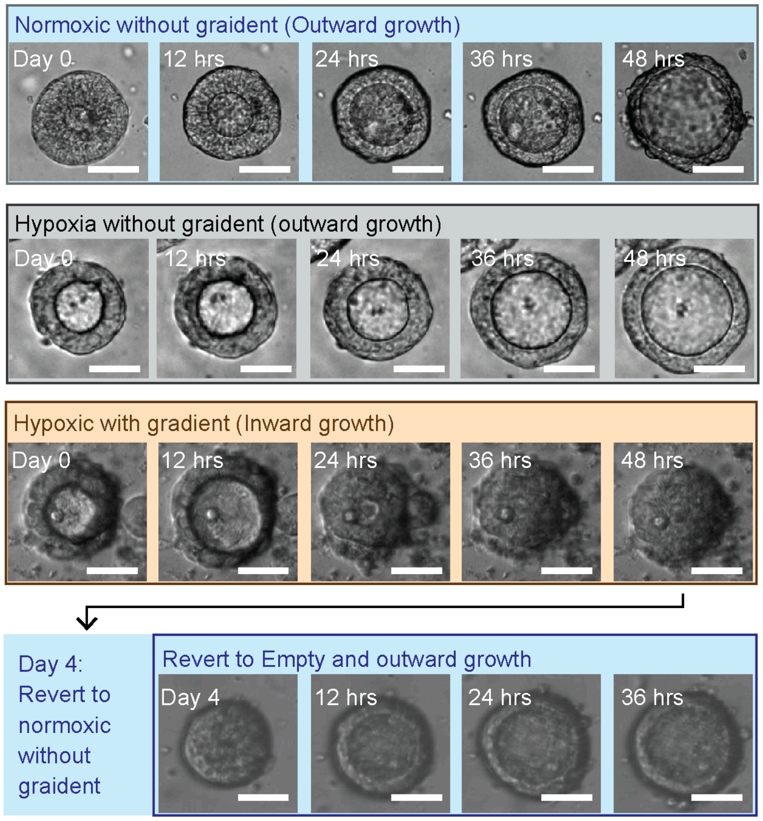


**Extended Data Fig. 7| Cancer organoids revert to cystic morphology after removal of oxygen gradient.**

### Methods

**Organoid samples.** Human organoid generation and culture was performed as previously described8. Briefly, patients undergoing colorectal resection and metastasectomy at MSKCC were identified by chart review. Informed consent for bio-specimen tissue collection of primary tumor was obtained through MSK IRB protocols 06-107, 12-245, 14-244 or 22-404. Freshly resected surgical tissue in surplus of clinical diagnostic requirements was processed for organoid generation. Human tumor organoids were maintained in Matrigel and cultured in HISC media (Advanced DMEM/F12 (AdDF12; Thermo Fisher Scientific), GlutaMAX (2 mM, Thermo Fisher Scientific), HEPES (10 mM, Thermo Fisher Scientific), N-acetyl-l-cysteine (1 mM, Sigma-Aldrich), B27 supplement with vitamin A (Thermo Fisher Scientific), Primocin (100 μg ml−1, InvivoGen), EGF (50 ng ml−1, Peprotech), Noggin (100 ng ml−1, Peprotech), A8301 (500 nM, Sigma-Aldrich), FGF2 (50 ng ml−1, Peprotech), IGF-I (100 ng ml−1, Peprotech). All animal procedures were approved by the Institutional Animal Care and Use Committee (IACUC) of Memorial Sloan Kettering Cancer Center. Normal mouse organoids were derived from C57BL/6J mice (strain 000664, Jackson Laboratories). Colons were dissected, flushed with PBS to remove faecal content, opened longitudinally, and cut into ~1 cm segments. Tissues were incubated in dissociation buffer (PBS with 8 mM EDTA, 0.5 mM DTT, and 10 U/mL DNase I (Roche 04716728001)) at 4C with gentle shaking for 60 minutes. Crypts were detached by vigorous shaking, pelleted, and further dissociated with TrypLE (Thermo Fisher Scientific). The resulting cell suspension was plated in and cultured in mouse HISC media (Advanced DMEM/F12 (AdDF12; Thermo Fisher Scientific), GlutaMAX (2 mM, Thermo Fisher Scientific), HEPES (10 mM, Thermo Fisher Scientific), N-acetyl-l-cysteine (1 mM, Sigma-Aldrich), B27 supplement with vitamin A (Thermo Fisher Scientific), N2 Supplement (ThermoFisher Scientific), Primocin (100 μg ml−1, InvivoGen), EGF (50 ng ml−1, Peprotech), murine Noggin (100 ng ml−1, Peprotech), A8301 (500 nM, Sigma-Aldrich), FGF2 (50 ng ml−1, Peprotech), IGF-I (100 ng ml−1, Peprotech),NGS-WNT (1 nM, ImmunePrecise N0001), and murine R-spondin 1 (1 μg mL-1, Peprotech). Established organoids were maintained in 6 well plates, dissociated to single cells (TrypLE, Thermo Fisher Scientific) for chemostat seeding. Organoids were routinely tested for mycoplasma contamination (MycoALERT PLUS detection kit, Lonza).

**Lentiviral generation and transduction.** Lentivirus was generated as previously described (PMID: 27246018). Briefly, pLenti-HRE-dUnaG (UnaG) was a gift from Roland Friedel (Addgene plasmid # 124372 ; http://n2t.net/addgene:124372 ; RRID:Addgene_124372) and pLV 4xSTAR-mScarletI-NLS-blast (STAR) was a gift from Hugo Snippert (Addgene plasmid # 136260 ; http://n2t.net/addgene:136260 ; RRID:Addgene_136260). HEK293T cells (ATCC) were cultured in DMEM supplemented with 10% FBS, GlutaMAX (2 mM, Thermo Fisher Scientific) and penicillin–streptomycin (100 IU ml−1, 0.1 mg ml−1, Thermo Fisher Scientific). HEK29T cells were transfected with 2nd generation packaging and envelope plasmids PAX (Addgene#12259) and MD2G (addgene), and UnaG or STAR plasmids. Media was replenished after 24 hours, and media containing virus was collected at 72 hours post transfection. Virus was concentrated with Lenti-X Concentrator (Takara Bio) overnight at 4C. Organoids were transduced with UnaG and STAR reporter virus, and antibiotic selection with puromycin or blasticidin, respectively, was initiated 72 hours after transduction. Organoid reporter lines were considered established after 3 passages. All cells were routinely tested for mycoplasma contamination (MycoALERT PLUS detection kit, Lonza).

**COMSOL simulation.** Numerical simulation was performed using finite element method (FEM)-based computational modeling software, COMSOL Multiphysics 5.4. Simulations were developed to solve the oxygen profiles in the fluidic cell-culture microchannel [16]. The COMSOL "Laminar Flow" interface with FEM formulations of the transport equation was applied to the 3D solution domain. This domain is a section of fluid in the channel (domain height: 0.25 mm, domain width: 1 mm). For the Materials selection, we used water, liquid to model the fluid inside channels, and PDMS to model the remaining PDMS structure. An "outlet" condition was applied to the two ends of the channel. And, the boundary conditions were set to "no slip" at the bottom and sides of the channel. The model was solved by a "stationary" solver to get the steady streaming pattern in the fluid domain.

**Microfluidic device fabrication.** The microfluidic device consisted of two parts: a microscope glass slide (75 mm × 25 mm × 1 mm, Fisher Scientific Inc.) and a PDMS channel layer with an embedded oxygen sensor.

The microfluidic channel was designed using AutoCAD and printed onto a photomask. The pattern was transferred onto a 4-inch silicon wafer via standard photolithography, using SU-8 2150 photoresist to produce microchannels with a height of approximately 200 µm. Polydimethylsiloxane (PDMS; Sylgard 184, Dow Corning) was prepared at a 10:1 base-to-curing-agent ratio and used to cast the device.

To integrate the oxygen sensor, a multilayer PDMS casting approach was employed. A small volume of PDMS was first poured onto the mold and spin-coated to form a ~250 µm layer, resulting in a ~50 µm PDMS layer above the channel ceiling after partial curing. At this stage, a pre-fabricated oxygen sensor strip (TROXSP5, PyroScience GmbH, Germany) was carefully positioned onto the sensing region. The sensor strip was trimmed into a 2 mm × 10 mm rectangle from a sensor patch originally supplied as a 10 mm × 10 mm square, and was aligned under a stereomicroscope to ensure consistent placement across devices.

After sensor placement, additional PDMS was poured to reach a total thickness of approximately 500 µm. The PDMS was then fully cured and peeled from the mold. Inlet and outlet ports were punched for tubing connections, and the PDMS layer was bonded to the glass slide using oxygen plasma treatment (AutoGlow 200, Glow Research)..

**Chemostat system fabrication.** The chemostat system included a 3D-printed scavenger tank mounted against the optical setup. Media perfusion is controlled by a VWR peristaltic pump. The scavenger tank is heated via water bath and hot plate with a safety shutoff at 37 °C to maintain stable temperatures. A custom LabVIEW program integrated control of the scanning stage, pump, light source, and imaging system. Brightfield imaging was performed using a VM-2 video microscope with integrated camera and 10X objective, enabling time-lapse capture of organoid cultures during oxygen gradient formation.

**Oxygen Sensing and Gradient Profiling.** Dissolved oxygen in the microfluidic chemostat device was measured using a contactless optical sensor system (PyroScience GmbH) comprising TROXSP5 sensor strips, a SPFIB-BARE optical fiber, and a FireSting-O2 meter. Sensor strips (2 mm × 10 mm) were embedded <200 µm from the organoid culture region and spanned the channel width. Fluorescence signals were converted to oxygen concentrations using Pyro Workbench software. Sensors were calibrated monthly using air-saturated water and 10 g/L sodium sulfite.

The optical fiber was mounted on a motorized XY stage (Thorlabs LTS150), controlled via LabVIEW, to scan oxygen along the Y-axis. Scanning parameters, including step size, number of steps, and dwell time, were customizable; typically, 22–23 positions were sampled at 0.4 mm intervals. At each location, 20 data points were collected over 20 seconds; the first and last 5 were discarded to minimize motion artifacts, and the middle 10 were averaged. Each scan took ~7 minutes, enabling continuous profiling (~7 scans/hour).

**Immunocyto staining and confocal imaging.** Organoids cultured under chemostat conditions were fixed *in situ* to preserve gradient-induced phenotypes. Briefly, live P146 organoids were incubated with 200 nM Mitotracker FM Deep Red (Thermo Fisher Scientific) in the chemostat for 30 minutes, washed twice with media, and then fixed with 4% paraformaldehyde in PBS for 30 minutes at room temperature. After fixation, samples were washed three times with PBS, permeabilized in 0.1% Triton X-100 in PBS for 10 minutes, and blocked in 5% bovine serum albumin (BSA) in PBS for 1 hour. F-actin was labeled by incubating with Alexa Fluor 488–conjugated phalloidin (Invitrogen; 1:200 in 1% BSA/PBS) and nuclei were counterstained with DAPI (1 µg $mL^{-1}$; Thermo Fisher Scientific). Following three PBS washes, the PDMS chips remained filled with PBS until imaging. All antibodies and dyes were diluted in 1% FCAs buffer, and wash steps consisted of gentle flushing with PBS. Whole-mount protocols (e.g., fixation in 4% PFA and permeabilization in Triton X-100) were adapted from established organoid immunofluorescence methods2[18]. Confocal images were acquired on either a Leica SP8 or a Nikon Ti inverted confocal microscope. Excitation/emission settings were as follows: DAPI, 405 nm/425–475 nm; Alexa 488 (phalloidin), 488 nm/500–550 nm; Mitotracker Deep Red, 640 nm/655–705 nm.

**Image processing.** All confocal datasets were processed and quantified in ImageJ. Z-stacks were imported as hyperstacks and subjected to a maximum-intensity projection. Lumen size was quantified by selecting enclosed regions of low F-actin signal or high contrast boundary in the bright field. The “Analyze” function was used to measure lumen area. Cell-covered area was determined by subtracting lumen area from the total cell size. All threshold levels and ROI settings were held constant across experimental groups. Quantification was performed from at least three independent chemostat runs.

**Patient pathology.** Archival formalin-fixed, paraffin-embedded (FFPE) clinical tissue blocks for immunostaining were identified by database search and chart review of patients who had signed pre-procedure informed consent to MSK IRB protocols 06-107, 12-245, 14-244 or 22-404 for biospecimen collection. Histopathological data interpretation were overseen by an expert gastrointestinal pathologist as previously described8.Tissue sectioning and hematoxylin and eosin (H&E) staining were performed by the MSKCC Molecular Cytology Core. For immunofluorescence (IF) staining, 5 um tissue sections were dewaxed (Histo-clear, cat# 50-899-90147), rehydrated, and steamed in antigen retrieva buffer (Abcam, ab93678) for 20 minutes. Cooled slides were washed twice with IF buffer (0.2% Triton X-10, 0.05% Tween, in PBS), blocked in 10% normal goat serum (Invitrogen, 50062Z) for 20 minutes, and incubated with primary mouse anti-TOMM20 antibody (1:100, Abcam, ab56783) diluted in 10% normal goat serum overnight at 4C. Following primary antibody incubation, slides were washed twice with IF buffer, incubated with TrueBlack™ Lipofuscin Autofluorescence Quencher (GoldBio, TB-250-1) diluted in 70% ethanol. After subsequent washes with 70% ethanol and IF buffer, slides were incubated with 594 anti-rabbit antibody (1:400, Invitrogen, A11012) for 1 hour at room temperature, washed three times with IF buffer and one time with cold DPBS. Slides were mounted using medium containing DAPI (Novus Biologics, H-1200-NB) and stored overnight at 4C prior to imaging on a Pannoramic Scanner (3DHistech, Budapest, Hungary) using a 40x/0.95NA objective.

**Statistical analysis.** The results are shown as mean ± standard error of the mean (SEM). To determine the statistical significance of the differences between the experimental groups, two-tailed unpaired Student’s t tests were performed using the Prism 10 software (GraphPad), as indicated in the figure. Sample sizes were based on experience and experimental complexity, but no methods were used to determine normal distribution of the samples. Differences reached significance with p values $< 0.05$ ($p$ values noted in figures). The figure captions contain the number of independent experiments or mice per group that were used in the respective experiments.

**Reporting Summary**
Further information on research design is available in the Nature Research Reporting Summary linked to this article.

**Data availability**
The authors declare that all data supporting the findings of this study are available within the paper and its Supplementary Information. Source data for the primary- sample figures are available.

**Code availability**
Custom LabView program and Matlab image processing code is available in the supplemental information.
